# Microbial Diversity of Co-occurring Heterotrophs in Cultures of Marine Picocyanobacteria

**DOI:** 10.1101/2020.07.15.205195

**Authors:** Sean M. Kearney, Elaina Thomas, Allison Coe, Sallie W. Chisholm

## Abstract

*Prochlorococcus* and *Synechococcus* are responsible for around 10% of global net primary productivity, serving as part of the foundation of marine food webs. Heterotrophic bacteria are often co-isolated with these picocyanobacteria in seawater enrichment cultures that contain no added organic carbon; heterotrophs grow on organic carbon supplied by the photolithoautotrophs. We have maintained these cultures of *Prochlorococcus* and *Synechococcus* for 100s to 1000s of generations; they represent ideal microcosms for examining the selective pressures shaping autotroph/heterotroph interactions. Here we examine the diversity of heterotrophs in 74 enrichment cultures of these picocyanobacteria obtained from diverse areas of the global oceans. Heterotroph community composition differed between clades and ecotypes of the autotrophic ‘hosts’ but there was significant overlap in heterotroph community composition. Differences were associated with timing, location, depth, and methods of isolation, suggesting the particular conditions surrounding isolation have a persistent effect on long-term culture composition. The majority of heterotrophs in the cultures are rare in the global ocean; enrichment conditions favor the opportunistic outgrowth of these rare bacteria. We did find a few examples, such as heterotrophs in the family Rhodobacteraceae, that are ubiquitous and abundant in cultures and in the global oceans; their abundance in the wild is also positively correlated with that of picocyanobacteria. Collectively, the cultures converged on similar compositions, likely from bottlenecking and selection that happens during the early stages of enrichment for the picocyanobacteria. We highlight the potential for examining ecologically relevant relationships by identifying patterns of distribution of culture-enriched organisms in the global oceans.

**IMPORTANCE:** One of the biggest challenges in marine microbial ecology is to begin to understand the rules that govern the self-assembly of these complex communities. The picocyanobacteria *Prochlorococcus* and *Synechococcus* comprise the most numerous photosynthetic organisms in the sea and supply a significant fraction of the organic carbon that feeds diverse heterotrophic microbes. When initially isolated into cultures, *Prochlorococcus* and *Synechococcus* carry with them select heterotrophic microorganisms that depend on them for organic carbon. The cultures self-assemble into stable communities of diverse microorganisms and are microcosms for understanding microbial interdependencies. Primarily faster-growing, relatively rare, copiotrophic heterotrophic bacteria – as opposed to oligotrophic bacteria that are abundant in picocyanobacterial habitats – are selected for in these cultures, suggesting that these copiotrophs experience these cultures as they would high carbon fluxes associated with particles, phycospheres of larger cells, or actual attachment to picocyanobacteria in the wild.

## INTRODUCTION

Microorganisms perform a variety of matter and energy transformations in the oceans that underlie global biogeochemical cycles. At the base of these transformations are photosynthesis and other forms of autotrophic fixation of carbon dioxide. The picocyanobacteria, *Prochlorococcus* and *Synechococcus*, contribute approximately 25% of this global ocean net primary productivity (Flombaum et al., 2013).

Abundant, free-living oligotrophic bacteria like *Prochlorococcus*, many strains of *Synechococcus*, and members of the SAR11 clade of heterotrophs exhibit genome streamlining, likely driven by reductive evolutionary pressures acting on bacteria that have large population sizes and live in relatively stable environments (Biller, Berube, et al., 2014; Giovannoni et al., 2005; Rocap et al., 2003). Together these three groups can represent more than 50% of free-living bacteria in the oligotrophic surface oceans (Becker et al., 2019; Giovannoni, 2017). By contrast, opportunistic marine copiotrophic organisms typically have larger genomes and cell sizes, and are subject to periodic fluctuations in abundance, typically depending on an influx of organic matter for growth (López-Pérez et al., 2012; Romera-Castillo et al., 2011; Tada et al., 2011; Zemb et al., 2010).

Unlike the low nutrient regimes of the oligotrophic oceans, cultures of *Prochlorococcus* and *Synechococcus* have high concentrations of inorganic nutrients (N, P, Fe) and accumulate dissolved organic matter and detritus as cells grow and die (Becker et al., 2014; Moore et al., 2007). These conditions might favor sympatric heterotrophic microorganisms adapted to higher organic carbon concentrations such as those associated with detrital particles or phytoplankton blooms (Buchan et al., 2014; Seymour et al., 2017; Stocker, 2012; Tada et al., 2011; Teeling et al., 2012). Even in oligotrophic oceans, the concentration of dissolved organic matter derived from phytoplankton can be hundreds of times higher in the diffusive boundary layer surrounding cells – the “phycosphere” (Bell & Mitchell, 1972; Seymour et al., 2017) than in the surrounding water. The size of the sphere is a function of the size of the cell producing it, thus large eukaryotic phytoplankton can create phycospheres extending 100s to 1000s of microns from their surface, while those of *Prochlorococcus* and *Synechococcus* span only a few cell lengths in diameter – i.e. 1-2 microns (Seymour et al., 2017). The assembly of heterotrophic bacterioplankton strongly depends on the nature of organic matter exudate (Amin et al., 2012; Durham et al., 2015, 2017; Fu et al., 2020; Gärdes et al., 2011; Goecke et al., 2013; Landa et al., 2017), which includes a diverse collection of compounds (Becker et al., 2014; Durham et al., 2015; Hellebust, 1967), and the bacteria in these communities can alter the physiology of their hosts (Amin et al., 2012; Durham et al., 2017).

While studies of the interactions between bacterioplankton in the phycosphere of eukaryotic hosts has received extensive attention, heterotrophs also exhibit chemotaxis towards metabolites derived from *Prochlorococcus* and *Synechococcus* (Seymour et al., 2010), hinting that some of the principles of interactions for large-size class phytoplankton may also hold for the picocyanobacteria. Instead of being derived from the relatively limited diffusive boundary layer around picocyanobacteria, free diffusion of dissolved organic carbon (DOC) in bulk seawater may be the most common way for heterotrophs in the oligotrophic oceans to obtain carbon for growth (Zehr et al., 2017). Hence, diffusion-limited uptake of DOC may limit growth for free-living heterotrophs in the oceans. Yet, there’s also evidence of direct interactions between picocyanobacteria and heterotrophs. Given the small size of picocyanobacteria, cells that attach directly to the picocyanobacterial host (apparent for some *Synechococcus* (Malfatti & Azam, 2009)) or associate with them in particulate matter (e.g. through heterotroph-mediated aggregation of cyanobacteria (Cruz & Neuer, 2019)) would be spatially co-localized enough to experience a so-called “phycosphere”.

Cyanobacterial cultures accumulate cells and organic carbon, which may lead to more frequent physical interactions as well as metabolic exchanges between the host cyanobacterium and associated heterotrophs, analogous to a phycosphere. Indeed, *Prochlorococcus* in culture exhibits both growth improvement and inhibition when grown in the presence of various co-occurring microbes (Aharonovich & Sher, 2016; Biller et al., 2016; Sher et al., 2011). The co-culture benefits include improved recovery from, and longevity in, stationary phase (Roth-Rosenberg et al., 2020), revival from low cell numbers during inoculation (Morris et al., 2008), expanded tolerance to temperature changes (Ma et al., 2017), and survival under extended darkness (Biller, Coe, et al., 2018; Coe et al., 2016). These various benefits have been attributed to the presence of these heterotrophic partners. However, culturing with heterotrophic bacteria can inhibit or delay the onset of growth in some strains of *Prochlorococcus* (Sher et al., 2011). Furthermore, at high concentrations *Alteromonas* strains that normally improve *Prochlorococcus* growth characteristics can inhibit it (Aharonovich & Sher, 2016).

When interactions are beneficial, heterotrophic bacteria often provide complementary functions for their “host” cyanobacteria. For example, *Prochlorococcus* strains lack the ability to produce catalase, which can detoxify radical oxygen species (ROS) formed during growth, but co-isolated heterotrophs that can produce catalase have been shown to complement this deficiency (Morris et al., 2008, 2011, 2012). The benefits provided to phototrophs by these heterotrophic microorganisms extend beyond removal of ROS, and include recycling and release of reduced N (López-Lozano et al., 2002; Zubkov et al., 2003) and P (Christie-Oleza et al., 2017) for resumption of autotrophic growth. In some cases, this nutrient recycling can allow cultures to have apparent indefinite longevity without transfer to fresh medium. *Synechococcus* can exhibit long-lasting growth when co-cultured with heterotrophic bacteria (Christie-Oleza et al., 2017), and we too have found that in co-cultures of *Prochlorococcus str.* MIT9313 with a *Rhizobium* species, and *Prochlorococcus str.* NATL2A with *Alteromonas macleodii* MIT1002, the cells can survive without transfer for over three years, in contrast to a few weeks for axenic cultures (unpublished data).

Because of their beneficial interdependencies, separating cyanobacteria from their heterotrophic “helpers” (*sensu* Morris et al., 2008) in enrichment cultures is a challenge when trying to obtain pure cultures (Christie-Oleza et al., 2017; Morris et al., 2008). At the same time, however, xenic cultures of these phototrophs present a unique opportunity to investigate the effects of selection on self-assembled mixed microbial communities, and ultimately, probe their interdependencies. While the boom-bust and high nutrient concentration conditions of batch culture do not mimic the conditions present in the open ocean (except for on particles or in phytoplankton blooms), these cultures represent a theater for extending the phycosphere of these cells throughout the culture volume. In these phycosphere analogues, we can observe what selective forces act on indigenous communities, and ultimately explore their mechanistic underpinnings. These data can inform future work on the dynamics of particle-attachment or aggregation among picocyanobacteria and associated heterotrophs. The robustness of these cultures may also provide insights into designing more efficient methods for obtaining and sustaining axenic strains of picocyanobacteria.

We have maintained cultures of *Prochlorococcus* [56] and *Synechococcus* [18], and accompanying heterotrophic “bycatch” for 100s to 1000s of generations in the laboratory by serial liquid transfer. We used this collection to address several questions about the nature of these phototroph-associated microbial communities. What is the composition and diversity of these heterotroph communities after many passages? How does the composition of our long-term cultures compare to that in cultures of picocyanobacteria or diatoms maintained for less time? Which factors best determine composition of the heterotrophic community: the method of isolation, the age of the culture, the inoculum from which the culture was derived (i.e. the specific time and place of isolation)? And finally, how does the composition of heterotrophs in culture compare to the communities in the waters from which the host cyanobacterium was isolated?

## RESULTS AND DISCUSSION

### The cyanobacterial culture collection

The cyanobacterial isolates used in the analysis span multiple clades of *Prochlorococcus* and *Synechococcus* obtained from many locations and times (between 1965 and 2013) in the global oceans (Figure 1A, Table S1). The isolates – most of which are from the oligotrophic oceans – have been maintained by serial transfer for many years, with cultures ranging in age from 6 to 55 years, and thus represent what we believe to be stable consortia of single algal isolates associated with diverse heterotrophic bacterial communities (see Materials and Methods, and Figure S1) (Figure 1B).

**Figure 1.**
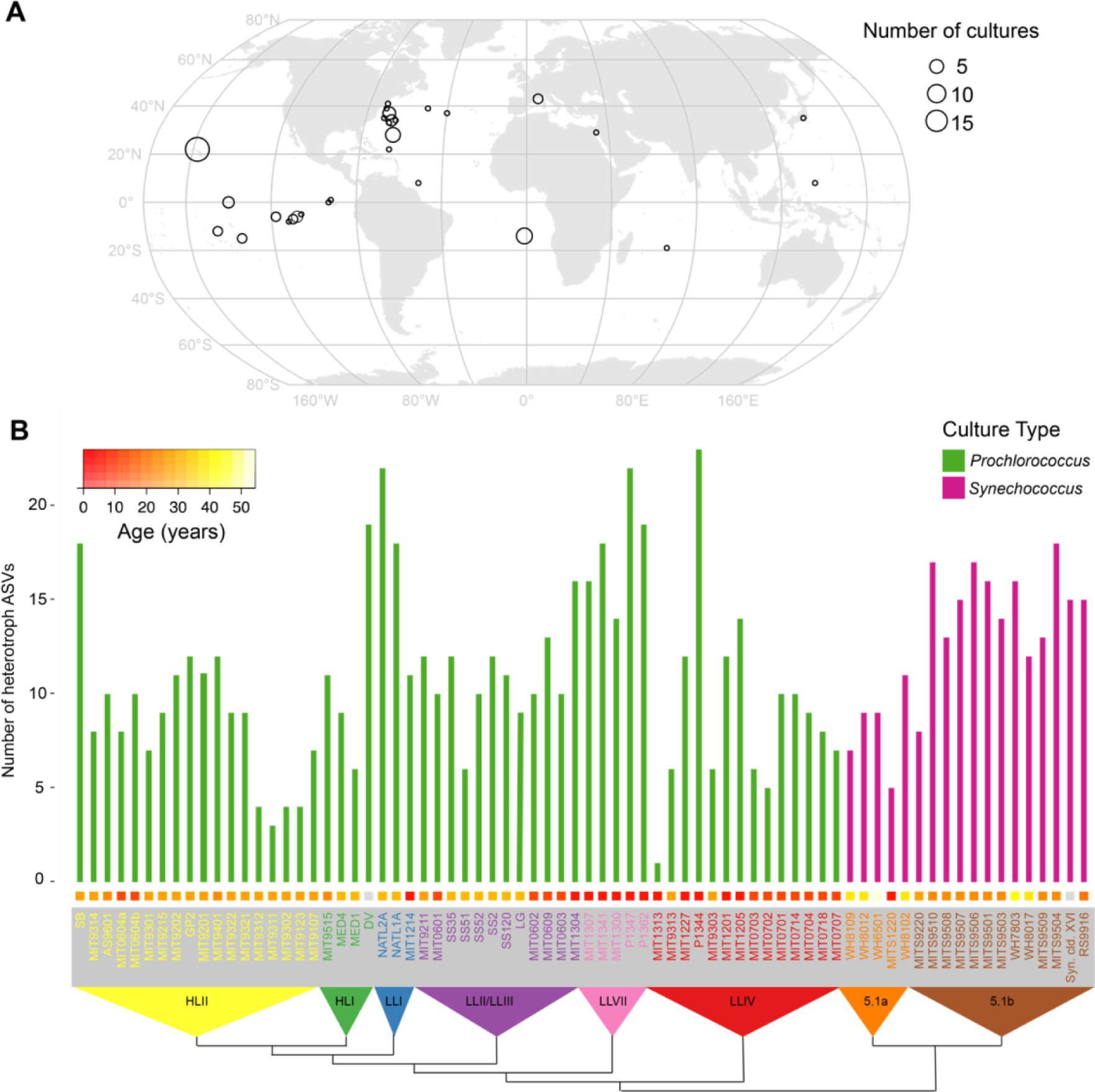
Microbial diversity in enrichment cultures from the global oceans. (A) Origin of *Prochlorococccus* and *Synechococcus* cultures obtained over several decades and maintained in serial transfer batch cultures for 100s – 1000s of generations. The size of each point on the map represents the number of cultures obtained from a given location. For date and methods of isolation see Table S1. (B) Richness of heterotrophic bacteria measured by the number of amplicon sequence variants (ASVs) exceeding a threshold relative abundance of 0.2% in *Prochlorococcus* (green) and *Synechococcus* (magenta) cultures. Richness is organized by the phylogeny of host organisms (built using the 16S-23S intertranscribed spacer (ITS) sequence, and collapsed to indicate monophyletic groups) indicated along the bottom. HL (high light) and LL (low light) designations in the triangles refer to light-adaptation features of *Prochlorococcus* ecotypes, as reviewed in (Biller et al., 2014), and 5.1a and 5.1b refer to subclusters of *Synechococcus* group 5.1 (Ahlgren & Rocap, 2012). Colored boxes above each culture name indicate the years between when the culture was isolated and this analysis, as specified by the color bar in the upper left-hand corner.

### Heterotroph communities in the cultures

We anticipated that heterotrophic culture richness – as defined by amplicon sequence variants (ASVs) of the V4 region of 16S rRNA gene – might decrease with age of the culture due to extinctions over time. However, the number of ASVs, which ranged from 1 to 23 (Figure 1B), was only weakly, and not significantly anti-correlated with the age of the culture (Spearman’s rho = -0.19, p-value = 0.11). NATL2A, for example, at nearly 30 years old, is one of the oldest cultures, but has 22 ASVs, while P1344 is 6 years old and contains 23 ASVs – a difference of just one ASV for an age difference of 24 years. Further, we sampled three sets of cultures (*Prochlorococcus* str. MED4, NATL2A, and MIT9313) in 2018 and one year later, and found a general correspondence in the community composition over time (Figure S1). Cultures of the same *Prochlorococcus* maintained by different individuals (MIT0604a & MIT0604b) also showed similar community composition, as well as cultures derived from the same starting enrichment culture (e.g. SS2, SS35, SS51, SS52, and SS120, were all derived from the LG culture (Table S1)) (Figure 2). These results suggest that the heterotroph communities in these cultures remain similar over time and independent maintenance, but exhibit slight drift over time (i.e. the cultures are similar but not identical in composition) (Figure S1).

**Figure 2.**
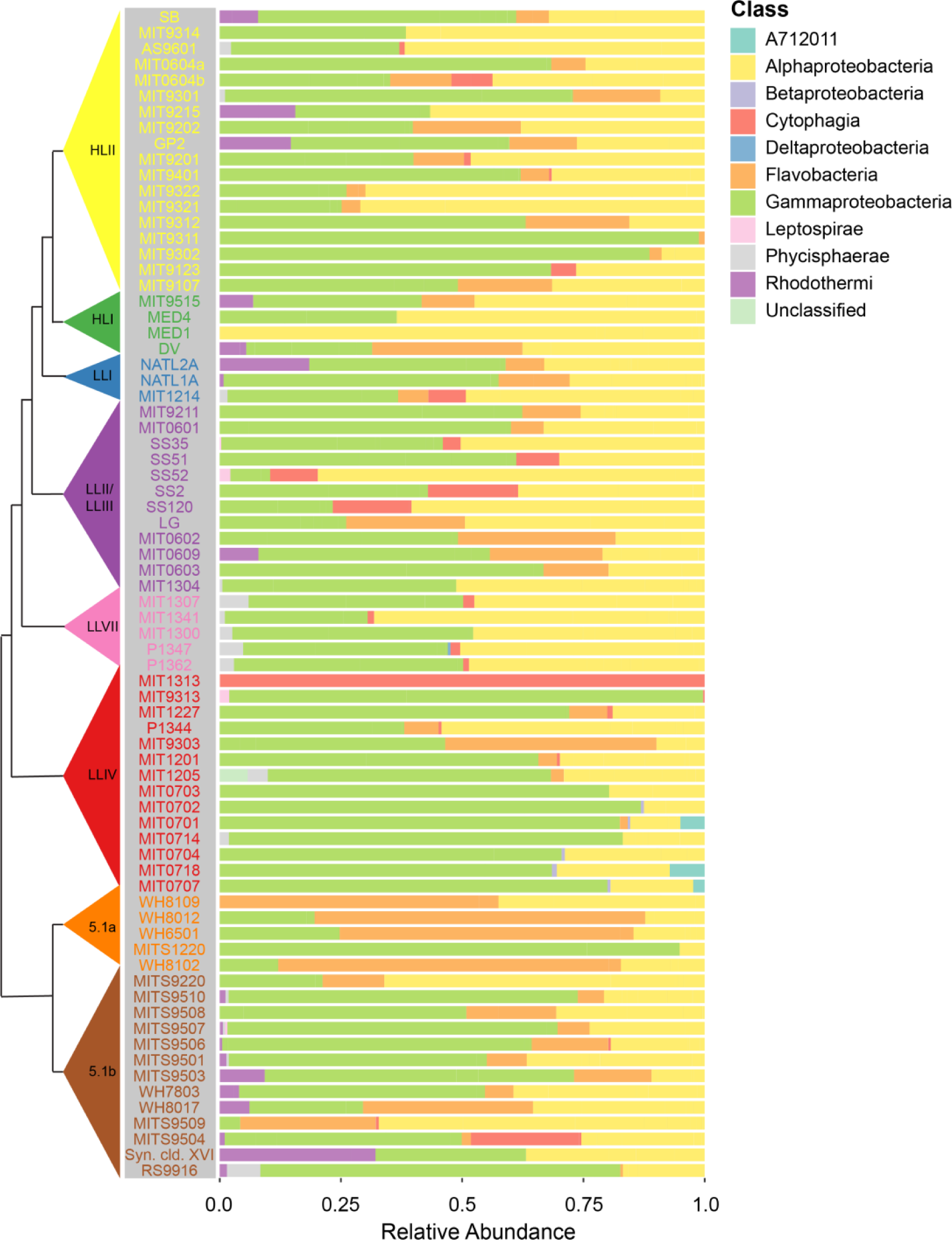
Composition of heterotroph communities, as defined by class membership of ASVs, in long-term cultures of *Prochlorococcus* and *Synechococcus* hosts (tree on the left as defined in Figure 1). The heterotroph communities in each culture are arranged using the phylogenetic tree of the host cyanobacterium based on the ITS sequence and collapsed into monophyletic groups as in Figure 1B. Relative abundance of heterotrophic community members, by class, is shown to the right of each strain name. To view the same data by hierarchical clustering see Figure S2.

We next explored whether the heterotrophic community composition was related to the cyanobacterial “host” – i.e. *Prochlorococcus* or *Synechococcus –* in the cultures. Which taxonomic groups were common to both, and which were more common in one or the other? Grouping ASVs at the phylum level, heterotroph communities in all cultures were largely comprised of bacteria from the phyla Proteobacteria (primarily in the classes Alpha- and Gammaproteobacteria, with some Delta- and Betaproteobacteria), Bacteroidetes (classes Cytophagia, Flavobacteria, and Rhodothermi), and Planctomycetes (class Phycisphaerae), with a minor contribution from Spirochaetes (class Leptospirae) and candidate phylum SBR1093 (class A712011) (Figure 2, Figure S2). SBR1093 was only in the LLIV clade (Figure 2). With respect to heterotrophic phyla, *Prochlorococcus* and *Synechococcus* cultures contain Proteobacteria and Planctomycetes at similar frequencies, but Bacteroidetes are more common, though not significantly, in *Synechococcus* cultures (94% versus 80%) (Figure 3A). At the class level, Alpha- and Gammaproteobacteria as well as Phycisphaerae have equal representation across the two cyanobacterial hosts (Figure 3B). Flavobacteria (90% of *Synechococcus* cultures versus 56% of *Prochlorococcus*, Fisher Exact Test p-value = 0.01) and Rhodothermi (56% of *Synechococcus* cultures versus 14% of *Prochlorococcus*, Fisher Exact Test p-value = 0.06), however, are more well-represented in *Synechococcus* cultures, suggesting that compounds in the exudate derived from *Synechococcus* may be better matched to their growth requirements. Indeed, *Prochlorococcus* and *Synechococcus* secrete distinct compounds during growth (Becker et al., 2014), few of which are characterized, but likely promote differential growth of associated heterotrophs as shown previously (Zheng et al., 2018, 2020).

**Figure 3.**
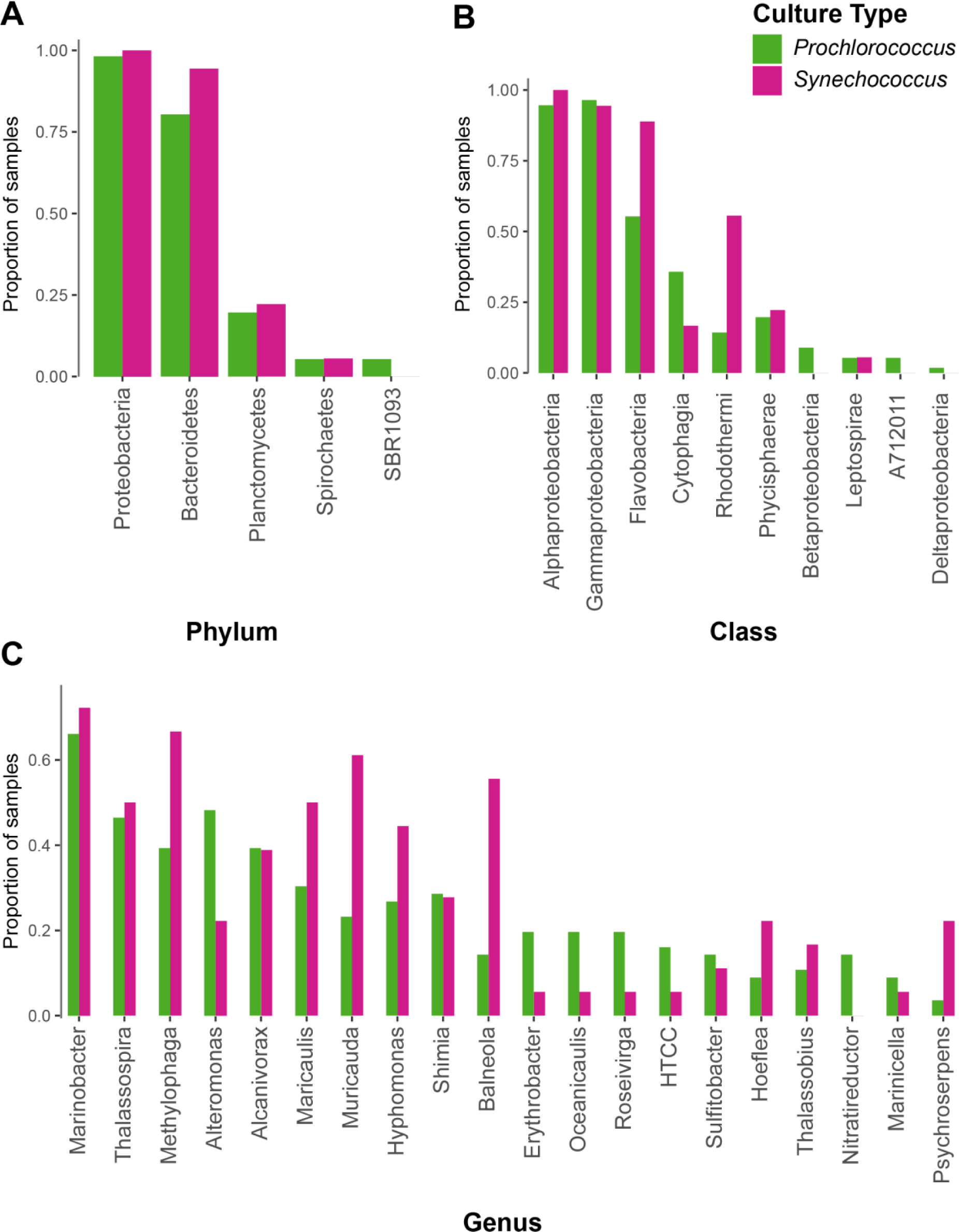
Proportion of cultures of either *Prochlorococcus* (green) or *Synechococcus* (magenta) containing ASVs belonging to a given bacterial (A) phylum, (B) class, or (C) genus (restricted to genera found in at least six cultures). Note that HTCC is listed in the Greengenes taxonomy, but is not formally recognized as a bacterial genus.

Minor differences in heterotroph communities between *Prochlorococcus* and *Synechococcus* hosts at the phylum and class level suggested that there might be more pronounced differences at the genus level. This was not the case, however; instead they were quite similar (Figure 3C). For instance, the most ubiquitous genus (present in over 60% of both culture types) is *Marinobacter* (Figure 3C), which is well-known to be associated with picocyanobacteria in culture (Morris et al., 2008). In addition to *Marinobacter*, other genera present in at least 15% of cultures including *Thalassospira, Methylophaga, Alteromonas, Alcanivorax, Maricaulis, Muricauda*, and *Hyphomonas* (but not *Shimia*) have been associated with metabolism of hydrocarbons or C1 compounds derived from lipid catabolism (Coulon et al., 2007; Dong et al., 2018; Hara et al., 2003; Kappell et al., 2014; Koch et al., 2020; Kostka et al., 2011; Lea-Smith et al., 2015; Liu et al., 2007; López-Pérez et al., 2012; Neufeld et al., 2007; Vila et al., 2010; Yakimov et al., 2007; Zhao et al., 2010), suggesting that hydrocarbon metabolism might play an important role in their growth in these cultures. Indeed, previous work showed an upregulation of genes for fatty acid metabolism including lipid beta-oxidation in co-culture of *Alteromonas macleodii* with *Prochlorococcus* (Biller et al., 2016*). Prochlorococcus* secretes vesicles (potentially a source of lipids) in culture and in the wild (Biller et al., 2017; Biller, Schubotz, et al., 2014), which can outnumber cells by a factor of 10 or more, and marine heterotrophs like *Alteromonas* are capable of growing on these vesicles as a sole source of carbon (Biller, Schubotz, et al., 2014). Recent work also suggests that *Alteromonas* isolates derived from cultures of *Prochlorococcus* carry genes for the degradation of aromatic compounds produced by the cyanobacterium (Koch et al., 2020), which may provide a selective advantage. Additionally, metagenomics on a *Synechococcus*-associated culture revealed high abundance of TonB-dependent transporters in *Muricauda*, potentially involved in lipid uptake, and proteins involved in the export of lipids in the proteome of the *Synechococcus* host (Zheng et al., 2020).

Finally, we investigated whether specific ASVs were differentially represented between culture types (*Prochlorococcus* vs *Synechococcus*). Using indicator species analysis (see Materials and Methods), a method to identify taxa associated with a specific habitat (here either a *Synechococcus* or *Prochlorococcus* culture), we found that 20 ASVs were more prevalent among *Synechococcus* cultures than *Prochlorococcus* cultures, but only one ASV (a *Marinobacter* sequence) more prevalent in *Prochlorococcus* cultures (Table S2). These associations strengthen the possibility that the structure of heterotrophic communities may arise in response to cyanobacterial host-specific secretion of compounds.

Consistent with there being more indicator species in *Synechococcus* cultures, community composition was more similar across *Synechococcus* cultures than *Prochlorococcus* cultures (as measured by unweighted UniFrac distance at the ASV level (PERMANOVA, p-value < 0.01)) (Figure 4A, Figures S3 & S4). Further, heterotroph community composition using the same metric was significantly associated (PERMANOVA, p-value < 0.01) with cyanobacterial ecotype (Figure S4A) and clade (Figure S4C), but there were no discernible patterns in these associations.

**Figure 4.**
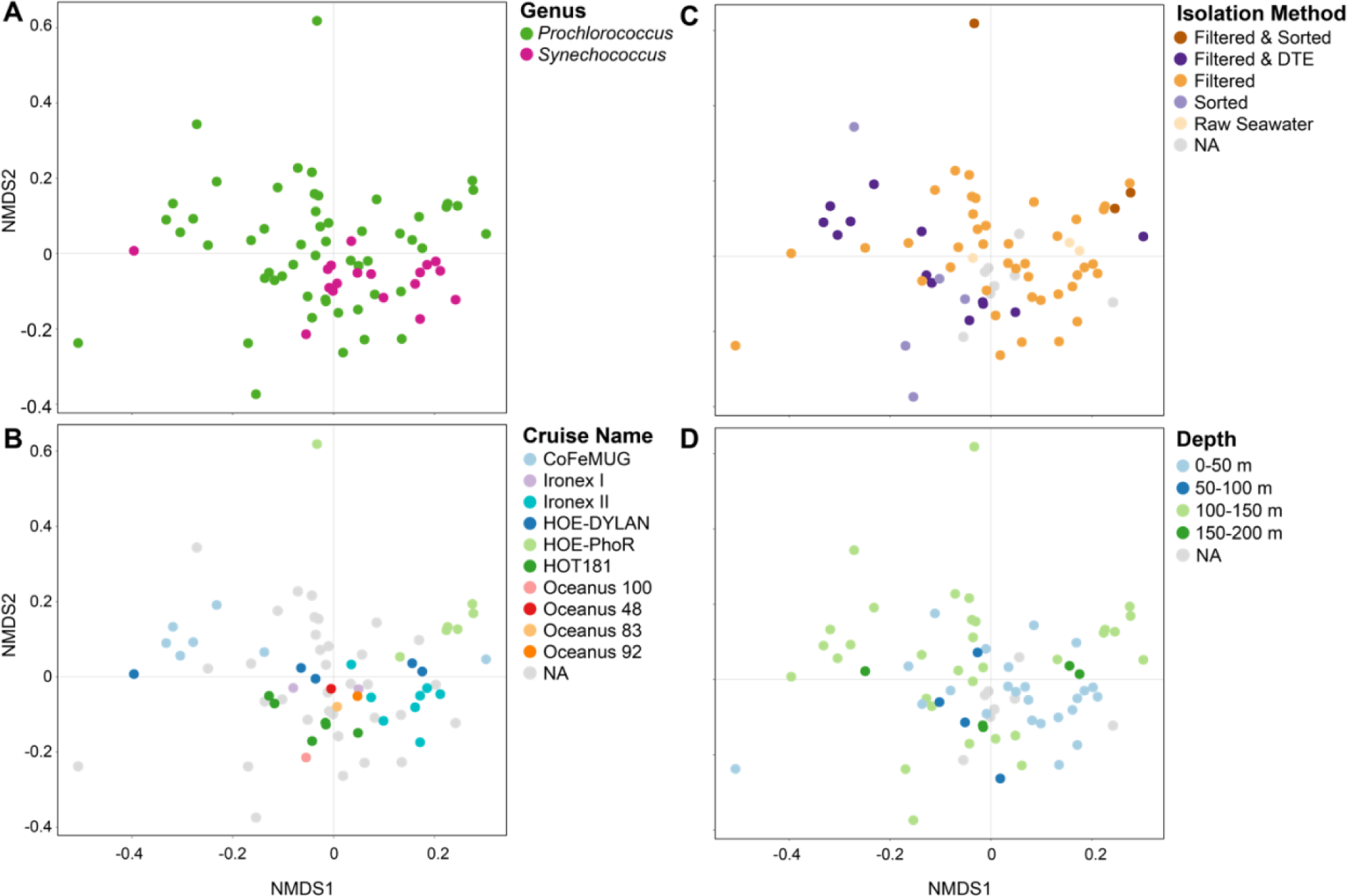
Ordination of heterotroph community composition at the ASV level overlaid with metadata pertaining to conditions of isolation of enrichment cultures. Non-metric multidimensional scaling (NMDS) of heterotroph communities using unweighted UniFrac as the distance metric overlaid with (A) the host cyanobacterium genus in the enrichment culture, (B) the cruise name from which the culture was isolated, (C) the isolation method (pre-filtered to exclude larger cells, pre-filtered and sorted via flow cytometry, pre-filtered and diluted-to-extinction (DTE), not filtered and sorted via flow cytometry, or cloned from a mother culture), and (D) the depth of the seawater sample. The closeness of two points in NMDS space reflects the distance between communities, with communities having more similar phylogenetic structure (as measured by unweighted UniFrac) grouping more closely together. Heterotroph communities for which metadata was not known are indicated as NAs. See also Figures S3, and S4 for relationships between community structure and ecotype, clade, isolation location, and culture age.

### Integrating the dataset with similar studies

To assess the similarity of other phytoplankton-associated microbiomes to those obtained in this study we re-processed - in the context of our dataset – 16S rDNA amplicon sequence data as 97% OTUs (see Materials and Methods) – from published diatom (Behringer et al., 2018) and recently isolated *Synechococcus*-associated microbial communities (Zheng et al., 2018). Surprisingly, the heterotrophic communities in diatom and *Synechococcus* cultures each shared almost half the OTUs found in our cultures: 45% (13/29) for the diatoms and 41% (29/70) for the *Synechococcus* cultures (Table S3, Figure S5). Among these OTUs, 11 were present in cultures of all three groups of phytoplankton, potentially comprising a ‘core’ set of sequence clusters associated with phytoplankton isolates. These include three of the ubiquitous genera (*Roseivirga, Maricaulis*, and *Alteromonas*) from this study, which derive from three separate classes (Figure 3C). At a high level, the taxonomic diversity in cultures from *A. tamarense* and *T. pseudonona* in a separate study (Fu et al., 2020) exhibited many of the same marine heterotrophs seen here (Figure S6), which suggests a generic selective effect of diverse phytoplankton on their associated microbial communities.

In contrast to this core of shared bacteria, several additional classes (Acidimicrobiia, Actinobacteria, Planctomycetes class OM190, and Saprospirae) were represented in the Zheng et al. 2018 *Synechococcus* dataset, but absent from our study; only the class Saprospirae was present in the diatom dataset and absent in ours (Figure S6). Notably, the Actinobacteria from the *Synechococcus* dataset were only found in isolates from eutrophic waters; by contrast, all except for two (*Prochlorococcus* SB & *Synechococcus* WH8017) of the isolates obtained in this study were from oligotrophic waters, while the diatoms of Behringer et al. 2018 were isolated from coastal waters. Further work should examine the extent to which these differences are driven by differences in starting inoculum versus physiological features of the host phytoplankton.

### Relationships between heterotroph community and features of the sample of origin

We examined the extent to which differences in heterotroph communities – here measured at the level of ASVs – in the cultures were related to factors involved in isolation (Table S1). Specifically, we investigated whether the heterotroph composition between all pairs of communities varied with: cruise on which it was isolated, isolation method, sample location and depth, and date the culture was isolated. Each of the tested metadata variables showed a significant association (PERMANOVA, p-value < 0.01) with differences in community composition as measured by unweighted UniFrac distance (a measure of the phylogenetic similarity between two communities), and there were no obvious cases in which the pattern of associations for one variable completely overlapped those for another (Figure 4, Figure S3, Figure S4). Notably, richness (number of ASVs) in the cultures showed no association with any of the tested variables, as previously mentioned for culture age (Wilcoxon Rank Sum Test, p-value > 0.05). In other words, differences in the heterotroph membership of communities (unweighted UniFrac distance) associated with features of the sample of origin did not arise from differences in the total number of heterotroph ASVs (richness).

While “cruise” itself is not a particularly informative variable on its own, we did see a reproducible tendency for cultures obtained on the same cruise to have similar heterotroph community composition. For example, cultures with names beginning in MIT13 or P13 (i.e. MIT13*XX* or P13*X*), were isolated by the same person on the same cruise (HOE-PhoR) in 2013 from a depth of 150 m, and include *Prochlorococcus* from multiple different low light clades (LLIV, LLII/LLIII, and LLVII) (Table S1). They were isolated with a variety of methods including filtration, flow sorting, and dilution-to-extinction. With two exceptions (the LLIV cultures P1344 and MIT1313, the latter of which was dominated by a single heterotroph ASV after flow sorting the *Prochlorococcus*) from the suite of eight cultures, the MIT13*XX* and P13*X* strains clustered together by unweighted UniFrac, suggesting a sensitive dependence of microbial community diversity on the conditions – not solely the physical isolation methodology – under which the culture was first isolated (Figure 4B). These conditions include the initial composition of heterotrophic bacteria in the collected seawater sample, the specific media formulation or light and temperature conditions used for a given enrichment attempt, and the duration of time an enrichment culture had been maintained before derivation of individual algal strains. Together, these factors may drive the observation that cultures obtained from a given cruise frequently share similar communities (seen also with the CoFeMUG and EqPac/IRONEX cruises) (Figure 4B). We think these similarities result from a bottlenecking of community diversity shortly after cultures are sampled and phytoplankton are enriched.

To look into the effects of potential bottlenecking during the initial culturing phase, we examined the relationship between culture composition and physical isolation methods (Figure 4C). We see a tendency for cultures obtained using only filtration (not accompanied by dilution-to-extinction or flow sorting) to be more similar in heterotroph composition to each other than cultures also subjected to dilution-to-extinction or flow sorting (Figure 4C). Again, however, communities tended to cluster more by the corresponding cruise than by isolation method. These findings suggest that stochastic loss of heterotrophs that might occur by dilution-to-extinction reduces the structural convergence of heterotroph communities. Finally, heterotroph community composition in the cultures differs by collection depth (Figure 4D). For example, cultures isolated from 0-50 m and 50-100 m tend to cluster more tightly with each other than cultures isolated from 100-150 m and 150-200 m, consistent with the idea that community composition of heterotrophs varies with depth (Agogué et al., 2011; DeLong et al., 2006; Giovannoni & Stingl, 2005). This finding is not surprising given that light and nutrient gradients in the water column lead to more complex biogeochemical regimes – and hence a gradient in dissolved organic carbon compounds (Hansell & Carlson, 2001b, 2001a) from the surface.

In conducting the above analyses, we note that some of these variables are not independent (e.g. because ecotype depends on clade; or because strains were only isolated from one depth or one ecotype was isolated on a given cruise), so associations may arise from interdependencies between factors (Figure S7). These interdependencies may lead to associations with multiple variables which could be explained by a single variable. However, the overall trends show a clear linkage between initial conditions of culture isolation and the long-term composition of the culture.

### Comparison of cultured communities to distributions of their members in the wild

Because all of the cultures are sourced from seawater samples, we wanted to determine how heterotrophs present in cultures were represented in the global oceans. We used the bioGEOTRACES metagenomics dataset, which spans 610 samples over time and at multiple depths in locations throughout the Atlantic and Pacific Oceans (Biller, Berube, et al., 2018) to examine the spatial distributions of heterotroph communities present in our cyanobacterial cultures (Figure 5). We obtained reads mapping to the V4 region of the 16S gene, and clustered the ASVs from our culture collection with the global oceans data at 97% similarity (i.e. 97% OTUs) to accommodate differences in the nature of the data and processing (see Materials and Methods for details). The heterotrophs that dominate global ocean datasets (primarily belonging to the Pelagibacteraceae within the Pelagibacterales) are not well represented in our cultures (Figure 5B, Figure S8). This absence is not surprising as simply culturing oligotrophs like SAR11 is extremely challenging (Button, 1991; Carini et al., 2013; Cho & Giovannoni, 2004); thus, we expected that the high nutrient concentrations and periodic dilution of cells from culture transfers would enrich for opportunistic copiotrophic organisms.

**Figure 5.**
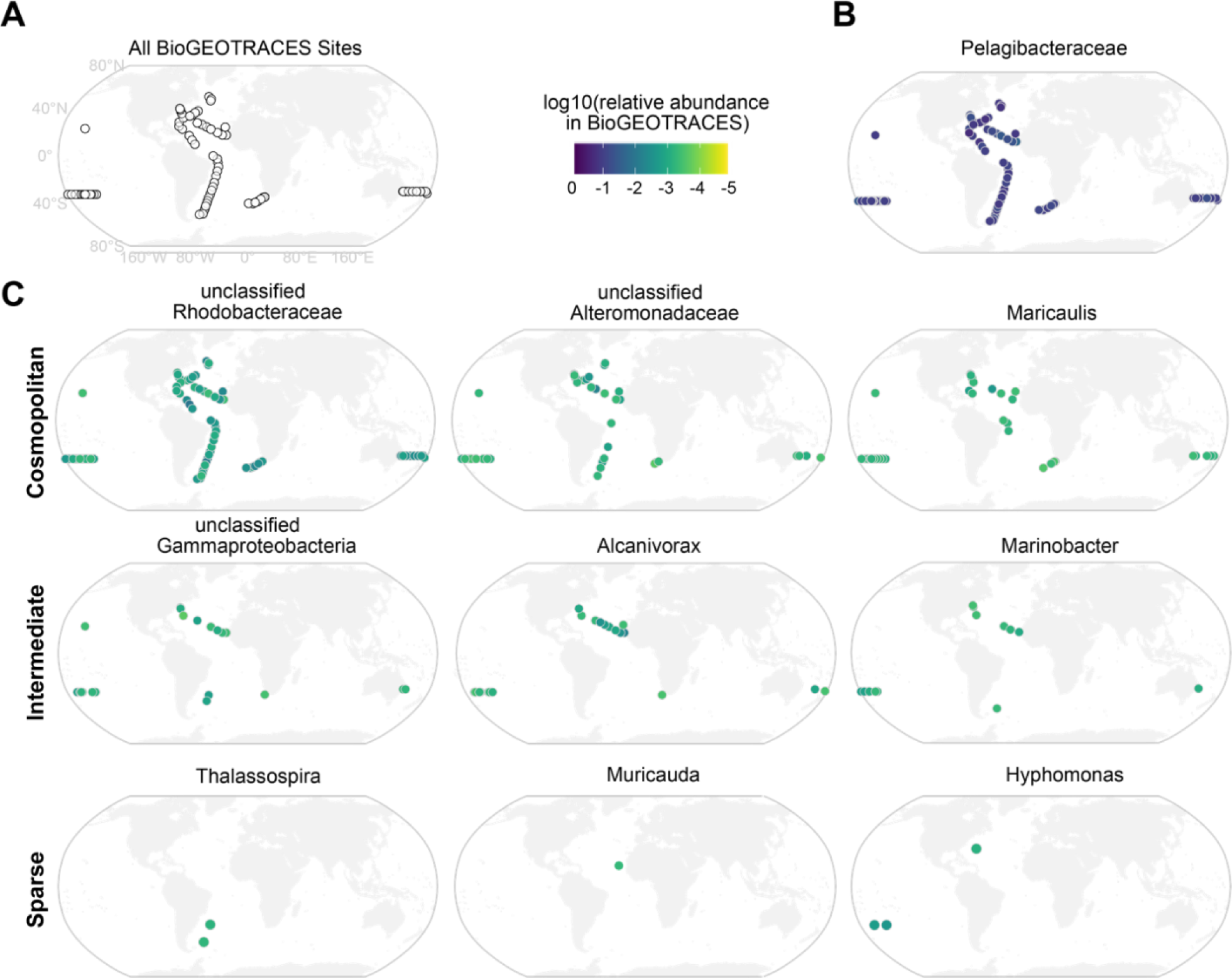
Representation of picocyanobacterial enrichment culture heterotrophic bacterial OTUs in global ocean surveys. (A) Location of the sampling sites of the bioGEOTRACES expeditions. (B) Example of a heterotrophic OTU (in the family Pelagibacteraceae) abundant in bioGEOTRACES, but absent from cultures. (C) Distribution of the most ubiquitous OTUs in cultures across the bioGEOTRACES sites. Such OTUs were present in almost all bioGEOTRACES sites (Cosmopolitan), some sites (Intermediate), or only a few sites (Sparse). Scale bar indicates the log10 relative abundance (number of reads normalized by number of non*-Prochlorococcus*, non-*Synechococcus* reads) at a given site. See also Figure S8 for the distribution of some OTUs abundant in bioGEOTRACES, but not prevalent in culture.

We next asked how well the prevalent OTUs in cultures were represented in the bioGEOTRACES database. While a few OTUs were readily detected at most sites (Figure 5C, Figure S8), this was not the general tendency; most that were prevalent in the cultures were rare or sparsely detectable in the wild (Figure 5C). Of the OTUs that dominated in culture, cosmopolitan OTUs included those classified as *Maricaulis*, Alteromonodaceae, and Rhodobacteraceae, while unclassified Gammaproteobacteria, *Alcanivorax*, and *Marinobacter* were intermediately distributed OTUs present in several sites. Finally, OTUs in the genera *Muricauda, Thalassospira*, and *Hyphomonas* were generally sparse or absent across the bioGEOTRACES sites (Figure 5C).

The sparsity of these organisms in the global oceans coupled with their prevalence in cultures of *Prochlorococcus* and *Synechococcus* suggests that they are selected for by the rich culture conditions – conditions that must be patchily distributed in the oceans. As an example, the genus *Muricauda* (family Flavobacteriaceae), which had an OTU well-represented in cultures, but sparsely detectable in bioGEOTRACES (Figure 5C), is known to be particle-associated, potentially specializing in degradation of high molecular weight organic compounds (Enke et al., 2019). Across all cultures in this study, more heterotroph OTUs in culture were shared with bioGEOTRACES samples taken below the epipelagic zone (>200 m) than above (0.3% of OTUs above versus 0.64% below, Fisher Exact Test p-value = 1e-4). The increase in OTUs shared with cultures at depth might be because of the reliance of heterotrophs on organic carbon as a source of energy in both systems.

Given the cosmopolitan distribution of some of the prevalent culture OTUs in bioGEOTRACES, we expected that they might be positively correlated with picocyanobacterial abundance along the transects. Indeed, we find that across the bioGEOTRACES sites, there is a strong relationship (Spearman’s rho = 0.302, p-value = 0) between the abundance of the Rhodobacteraceae OTU, which is ubiquitous in cultures, and the combined abundance of *Prochlorococcus* and *Synechococcus*. Notably, none of the other prevalent culture OTUs showed this relationship, suggesting that this OTU in particular may be coupled to the dynamics of these cyanobacteria in oceans (Table S4). Further laboratory investigations of the interactions of these ubiquitous bacteria with marine picocyanobacteria should reveal interesting and relevant exchanges of matter and energy in the global oceans.

## CONCLUSIONS AND FUTURE DIRECTIONS

Although the culture collection being analyzed here was not designed with this study in mind, there are some generalizations we can extract from the heterotrophic “bycatch” that was selected for in the enrichment cultures, which may help guide experiments designed to unravel the co-dependencies in these micro-communities. First, as expected from their obligate oligotrophy, the heterotrophs that numerically dominate the open ocean habitats are not present in these xenic cultures. For most of them, including the abundant SAR11 group, high organic carbon conditions present an obstacle (Button, 1991; Cho & Giovannoni, 2004) and hence developing a defined medium for their growth was a challenge (Carini et al., 2013). Although we designed a medium that sustains co-cultures of SAR11 and *Prochlorococcus* in log phase (Becker et al., 2019), SAR11 dies precipitously when *Prochlorococcus* enters stationary phase, suggesting that strict oligotrophs may not be able to tolerate the accumulation of substrates that occur during *Prochlorococcus* growth – a feature that would have eliminated them in the initial stages of isolating the cyanobacterial strains in our culture collection. This challenge during stationary phase might derive from deleterious effects of high nutrient concentrations on streamlined cellular physiology, with oligotrophs unable to regulate transport rates in the face of high nutrient concentrations (Braakman et al., 2017).

The absence of oligotrophs in the enrichment cultures is in striking contrast to the presence of the numerous copiotrophic heterotrophic strains that thrive in these cultures (this study, and (Aharonovich & Sher, 2016, 2016; Biller, Coe, et al., 2018; Biller et al., 2016; Morris et al., 2008; Sher et al., 2011)) and even “bloom” when *Prochlorococcus* reaches stationary phase (Becker et al., 2019). Studies of heterotrophic community dynamics in cultures over the course of the exponential and sustained stationary-phase growth (Zheng et al., 2020) have revealed shifts in heterotrophic abundances according to their differential capacities to utilize high- and low-molecular weight dissolved organic compounds. Further work as in that study will enable detailed linkage between functional genes in heterotrophic bacteria for metabolism of cyanobacteria-derived photosynthate and the dynamics of these functions in the global oceans.

Interestingly, most of the heterotrophic strains that are widespread among the cultures are not very abundant in the wild. These copiotrophs appear to thrive on high concentrations of organic compounds experienced in culture – an environment that must be patchily distributed in the open ocean habitat (Stocker, 2012). While it is likely that individual phytoplankton can selectively permit the growth of particular heterotrophic bacteria, the sharing of over 40% of OTUs between our dataset and each of the other phytoplankton datasets explored here suggests that phytoplankton may modify their environments in similar ways that lead to conservation of bacterial groups across cultures. It is possible that ubiquitous heterotrophic bacteria in cultures are similar to “broad-range taxa” that process simple metabolic intermediates (Enke et al., 2019), and thus grow on compounds that are released as generic byproducts of phytoplankton growth. Indeed, such compounds are likely abundant on nutrient rich particles, or ephemeral patches of high organic carbon in the phycosphere of larger phytoplankton (Seymour et al., 2017; Stocker, 2012; Thornton, 2014).

Clearly, to understand the metabolic exchanges between picocyanobacteria and oligotrophic heterotrophs we will have to isolate sympatric strains from the same location. In contrast to standard enrichment approaches, which appear to favor the growth of fast-growing, phycosphere-enriched bacteria, using dilution-to-extinction techniques and low nutrient media is likely to yield more of the abundant, free-living bacteria characteristic of the oligotrophic oceans. Like all challenges with these microorganisms, it is only a matter of time and effort.

## MATERIALS AND METHODS

### Culture maintenance

Previously isolated *Synechococcus* and *Prochlorococcus* from sites spanning the global oceans (Table S1) are maintained by serial passage in natural seawater medium amended with Pro99 nutrients (800 uM N, 50 uM P, trace metals, no added carbon) under low light intensity (∼10 µmol photons m^-2^ s^-1^) in continuous light or a 13:11 light:dark incubator (Moore et al., 2007). Prior to DNA extractions, cultures were grown until mid-exponential growth phase to increase the comparability of the communities across all cultures.

### Describing the cyanobacteria-associated microbial community composition

We used amplicon sequence variants (hereafter ASVs) of 16S rDNA as our marker. We analyzed the composition of the community of heterotrophs in our isolates when harvested in mid-exponential growth. To this end, we inoculated maintenance cultures (n=74) into fresh medium and allowed them to grow until mid-exponential growth before collecting cells by centrifugation for preparation of 16S rDNA libraries. Sequence libraries had an average depth of 113,602 (min: 72,369) reads after quality filtering, and in most samples, reads derived predominately from the cyanobacterial host (*Synechococcus* or *Prochlorococcus*). For the purposes of characterizing the associated community, we excluded reads derived from *Prochlorococcus* or *Synechococcus* in downstream analyses. The cultures contained a median of 11 ASVs (range 1-23) that were at or above an abundance of 0.2%, and a median of 16 ASVs estimated by rarefaction (range 3-31) (Figure 1B). We do not make quantitative claims about the abundances of ASVs in samples because of the dependence of heterotroph abundance on cyanobacterial growth state (Becker et al., 2019; Zheng et al., 2018) and biases in amplicon sequence data (Polz & Cavanaugh, 1998); we instead focus on the distribution of observed ASVs, and higher order taxonomic classifications across the cultures.

### Sequence generation and processing

DNA samples were prepared by pelleting 5 mL of exponentially growing cultures by centrifugation at 8,000 x g for 15 min. DNA was extracted using the DNeasy Blood & Tissue Kit (QIAGEN Cat No. 69504) following manufacturer instructions. The V4 region of the 16S rRNA gene was amplified using the 515F-Y (5’-GTGYCAGCMGCCGCGGTAA-3’) and 926R (5’-CCGYCAATTYMTTTRAGTT-3’) primers (Parada et al., 2016), and sequencing libraries prepared in duplicate before pooling using a two-step protocol described previously (25), with the exception of the MED4, NATL2A, and MIT9313 cultures from 2018, which used the same two-step protocol, but were generated in a different sequencing run using 515F (5’-GTGCCAGCMGCCGCGGTAA-3’) and 806R (5’-GGACTACHVGGGTWTCTAAT-3’) primers (Preheim et al., 2013). The libraries were sequenced on an Illumina MiSeq (Illumina, San Diego, CA, USA) platform, using 250 bp paired-end reads (except for the 2018 MED4, NATL2A, and MIT9313 cultures, which used 150 bp paired-end reads). The fastq sequence data files were processed to generate a table of amplicon sequence variants (ASVs) using DADA2 in R (Callahan et al., 2016), with the parameters for each dataset specified in the corresponding R script (https://github.com/microbetrainer/Heterotrophs).

### Mock community construction for sequencing validation

We extracted DNA as described previously from eleven bacteria with distinct V4 16S regions: axenic cultures of *Prochlorococcus* strains MED4, NATL2A, MIT9313, and MIT9312; *Synechococcus* strains WH7803, and WH8102; *Polaribacter* MED152, *Alteromonas macleodii* MIT1002, and three other heterotrophic bacteria. We amplified the full 16S gene using the primers 27F 5’-AGAGTTTGATCMTGGCTCAG-3’ and 1492R 5’-GGTTACCTTGTTACGACTT-3’, verified the 16S sequence by Sanger sequencing (Eton Bioscience, Inc. San Diego, CA, USA), then cloned the 16S gene from each culture using the Zero Blunt™ TOPO™ PCR Cloning Kit (ThermoFisher Cat #450245). The resulting plasmids were combined either in equimolar concentrations or in a 2-fold dilution series (with the most concentrated 16S 1024x more concentrated than the least) with three technical replicates, which were included as samples in 16S DNA library preparation.

### 16S amplicon sequence data analysis

In order to limit the extent to which we were measuring contaminating PCR products from adjacent wells, we excluded ASVs that had anomalously low abundance in one well given its abundance in an adjacent well. Because this approach is conservative (and likely removing true positive ASVs), it will have the tendency to deflate the estimates of taxonomic richness within a sample. We applied the following filter: if the relative abundance of an ASV in a specified well was less than one-tenth that of the maximum relative abundance in an adjacent well on the 96-well plate, the relative abundance of the ASV in the specified well was set to 0. Based on analysis of the mock community data, we found no contaminating sequences at a relative abundance of more than 0.002, thus, sequences were excluded from a sample if they had a relative abundance lower than 0.002. This cutoff tends to be conservative, as rarefaction analysis retained more ASVs per sample overall. For all analyses not involving OTU clustering, the sequences were trimmed to the same length. The sequences were imported to Qiime2 (Bolyen et al., 2019), where they were de-replicated using VSEARCH (Rognes et al., 2016) and assigned taxonomies using the gg-13-8-99-nb-classifier (Caporaso et al., 2010; DeSantis et al., 2006). Using fasttree (Price et al., 2010) and mafft alignment (Katoh & Standley, 2013) within Qiime2, phylogeny of the non*-Prochlorococcus*, non-*Synechococcus* ASVs was created.

### *Comparative analysis of diatom &* Synechococcus *culture datasets*

We compared sequences identified in this study to sequences found in newly isolated *Synechococcus*. cultures (Zheng et al., 2018) and sequences isolated from the phycosphere of diatoms in culture (Behringer et al., 2018). The *Synechococcus* study (Zheng et al., 2018) used 520F (5′-AYTGGGYDTAAAGNG-3′) and 802R (5′-TACNVGGGTATCTAATCC-3′) for 16S data generation and the second study uses 515F (5′-GTGYCAGCMGCCGCGGTAA-3′) and 786R (5′-GGACTACNVGGGTWTCTAAT-3′). Both sets of primers produce sequences that are nested inside those generated in this study (515Y-F, 926R). The diatom dataset contained microbial communities measured over time in cultures of four strains of *Asterionellopsis glacialis* and three strains of *Nitzschia longissima*. We included the initial measurements of microbial communities from nine new *Synechococcus* isolates, four from oligotrophic waters and five from eutrophic waters. For this analysis, we clustered at 97% identity (hereafter, OTUs) due to differences in the generation of the amplicon data in the published studies. The sequences from both studies were processed in DADA2 to produce a table of ASVs, with the parameters specified in the corresponding R script (https://github.com/microbetrainer/Heterotrophs). We did not trim ASVs from these three datasets to the same length. We excluded ASVs that were identified as either mitochondria or chloroplast from analysis, and excluded ASVs from a sample if they had a relative abundance less than 0.002. In Qiime2 (Bolyen et al., 2019), the ASVs were clustered at 97% identity using VSEARCH (Rognes et al., 2016), and the taxonomy of each cluster was assigned using the gg-13-8-99-nb-classifier (Caporaso et al., 2010; DeSantis et al., 2006). The samples were clustered using Ward’s method of hierarchical clustering on the unweighted UniFrac distance at the 97% OTU level (Lozupone & Knight, 2005) calculated with the phyloseq package in R (McMurdie & Holmes, 2013).

### BioGEOTRACES comparisons

Reads mapping to the V4 region of the 16S gene between the 515F-Y and 926R priming regions were obtained from bioGEOTRACES metagenomes (Biller, Berube, et al., 2018) courtesy of Jesse McNichol (data and code can be found at https://osf.io/n5ftw/). Without trimming the ASVs to the same length, the bioGEOTRACES and culture ASVs were imported to Qiime2. Within Qiime2, the sequences were dereplicated and clustered at 97% identity with VSEARCH. The OTUs were then assigned taxonomies using the gg-13-8-99-nb-classifier and represented on maps using the accompanying bioGEOTRACES metadata.

### Comparison of 2018 and 2019 heterotroph communities

The cultures described in this study were sampled in the summer of 2019. The non-axenic cultures of *Prochlorococcus* MED4, MIT9313, and NATL2A were sampled in 2018 and again in 2019. To investigate whether the composition of heterotrophs changed over this period, we compared the ASVs in these cultures sampled over time. Using DADA2, we trimmed the primers (the first 20 bases) from the reads from the 2018 samples, as quality decreased at this point in the sequences. We trimmed the ASVs from the two sources to the same length. For both datasets, ASVs were excluded from a sample if they had a relative abundance less than 0.002. In Qiime2, the ASVs were de-replicated across the two datasets using VSEARCH and assigned taxonomies using the gg-13-8-99-nb-classifier. Within Qiime2, fasttree, and mafft were used to create a phylogeny of the non*-Prochlorococcus*, non*-Synechococcus* ASVs. The abundance of each ASV in each sample was normalized by the number of *Prochlorococcus* sequences in the sample. The Spearman correlation between each pair of samples was calculated based on the abundance of each non-*Prochlorococcus* ASV normalized by the number of *Prochlorococcus* reads.

### Accession numbers

The 16S rDNA sequence data from this study were deposited in the NCBI Sequence Read Archive under BioProject ID PRJNA607777.

## ACKNOWLEDGMENTS

This work was funded by grants from the Simons Foundation (Life Sciences Project Award ID 337262, S.W.C.; SCOPE Award ID 329108, S.W.C.). S.M.K is funded by the Simons Foundation Fellowship in Marine Microbial Ecology. Jesse McNichol graciously shared the data for reads mapping to the V4 region of the 16S gene from bioGEOTRACES and HOT and BATS. We thank Dave VanInsberghe and the Polz lab for the 16S amplicon sequencing of the three *Prochlorococcus* cultures from 2018. We thank everyone who has contributed to this work through the isolation and maintenance of *Prochlorococcus* and *Synechococcus* cultures. Specific contributors are listed as “Isolators” in Table S1. We thank Fatima Hussain, Paul Berube, and Nikolai Radzinski for critical reading of the manuscript.

## CRediT taxonomy for author contributions

Conceptualization, S.M.K., A.C., S.W.C.; Methodology, S.M.K., E.T., A.C.; Software, S.M.K., E.T.; Validation, S.M.K., E.T., A.C.; Formal Analysis, S.M.K., E.T.; Investigation, S.M.K., E.T., A.C.; Resources, S.M.K., A.C., S.W.C; Data Curation, S.M.K., E.T.; Writing – Original Draft, S.M.K.; Writing – Review & Editing, S.M.K., E.T., A.C., S.W.C; Visualization, S.M.K., E.T.; Supervision, S.M.K., A.C; Project Administration S.M.K., A.C.; Funding Acquisition, S.M.K., S.W.C.

## SUPPLEMENTARY FIGURES

**Figure S1.**
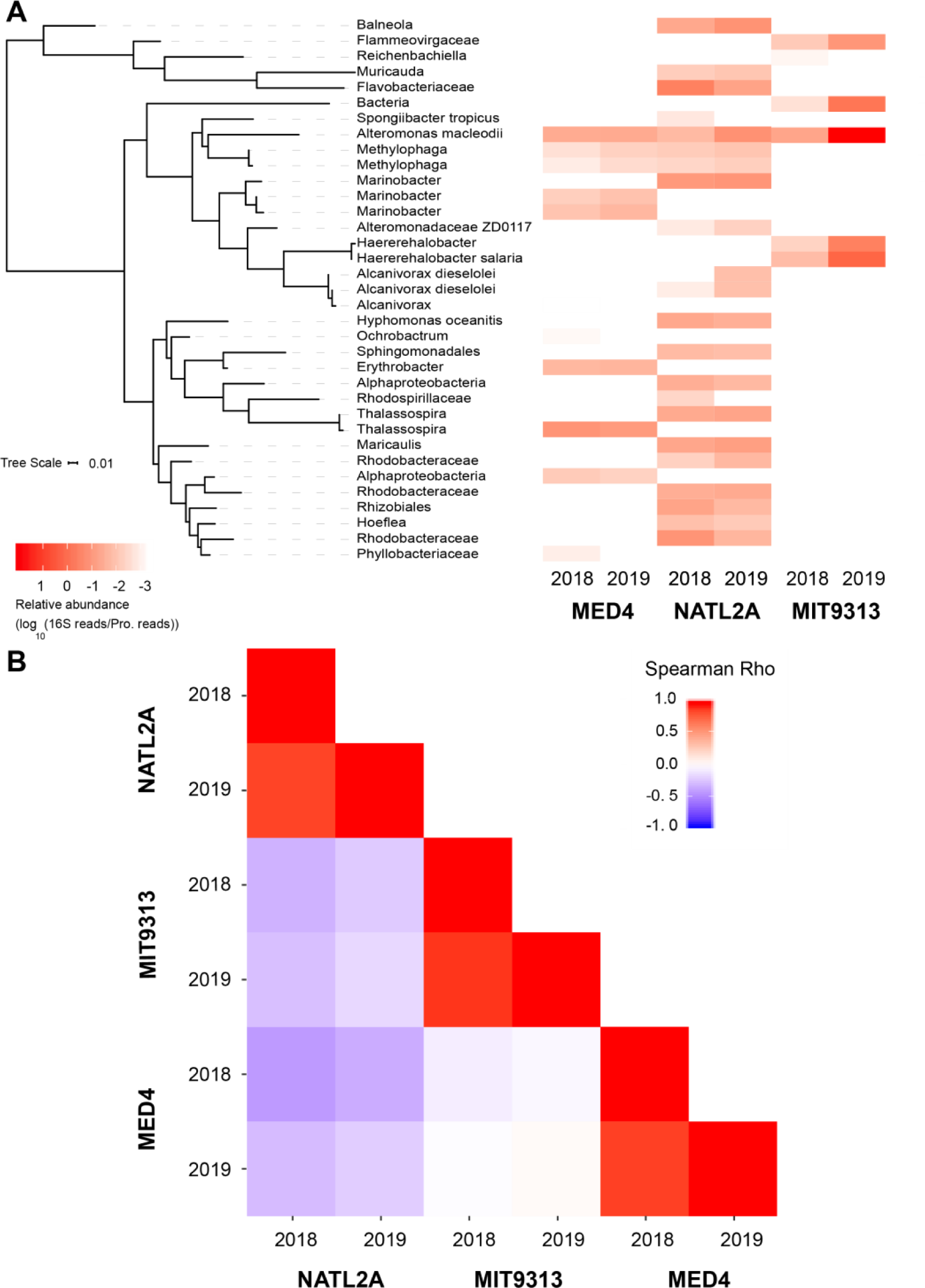
Stability of heterotroph community composition in three cultures sampled one year apart. (A) The *Prochlorococcus*-normalized relative abundance (indicated by the color bar) of each of the heterotroph ASVs is shown in the heatmap for each of three cyanobacterial enrichment cultures (*Prochlorococcus* strains MED4, MIT9313, and NATL2A) sampled once in 2018 and again in 2019 for amplicon sequencing. The phylogenetic tree on the left is based on the heterotroph ASV sequences. (B) The Spearman correlation (comparing the correspondence in rank) of relative abundance of heterotroph ASVs is indicated by the color bar.

**Figure S2.**
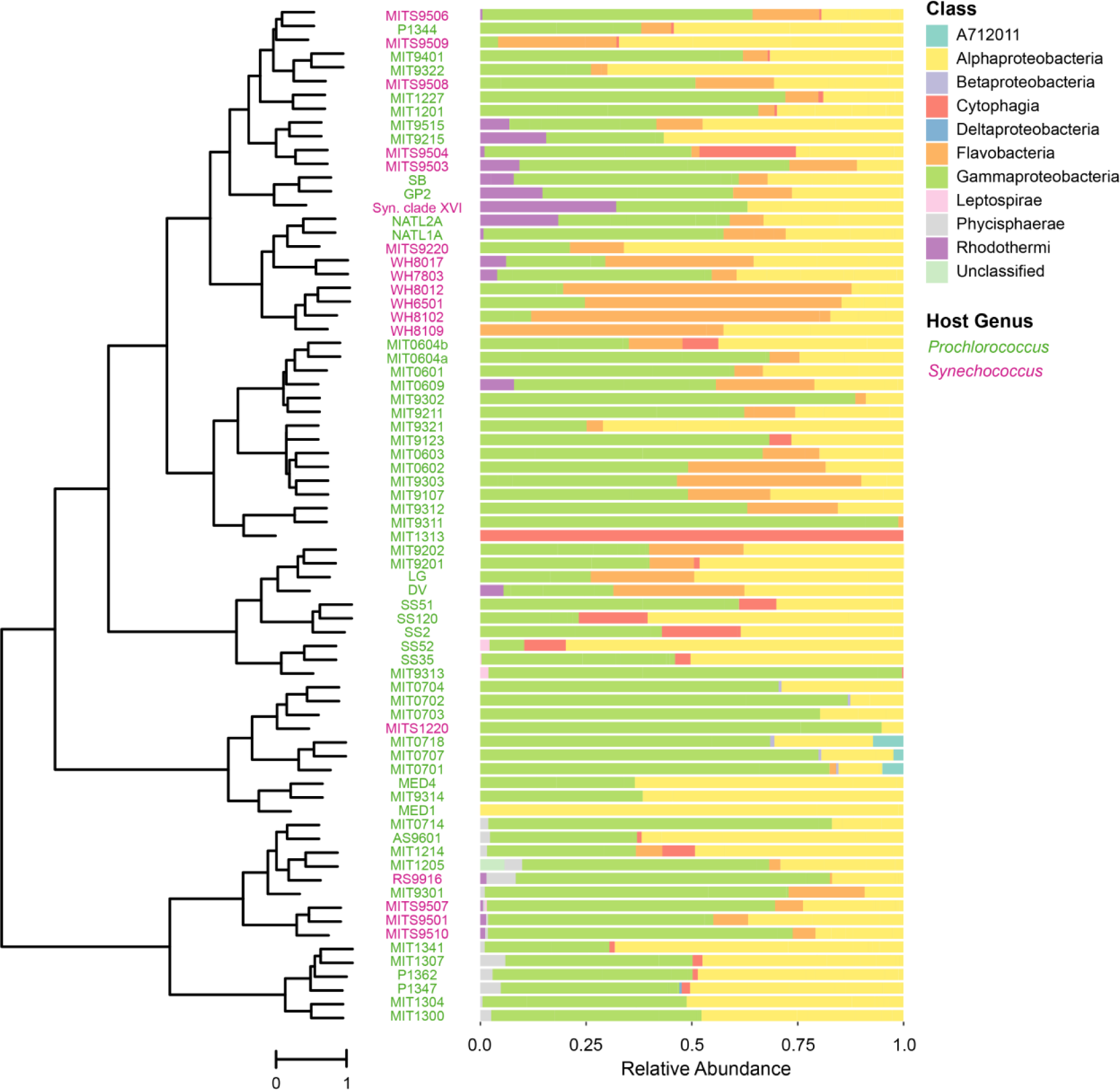
Composition of heterotroph communities, as defined by class membership of ASVs, in long-term cultures of *Prochlorococcus* (green) and *Synechococcus* (magenta) hosts. This is the same data as depicted in Fig. 2, but here the heterotroph communities in each culture are arranged using Ward’s method of hierarchical clustering with unweighted UniFrac on ASVs as the distance metric to emphasize the relationships between communities. Relative abundance of heterotrophic community members, by class, is shown to the right of each strain name.

**Figure S3.**
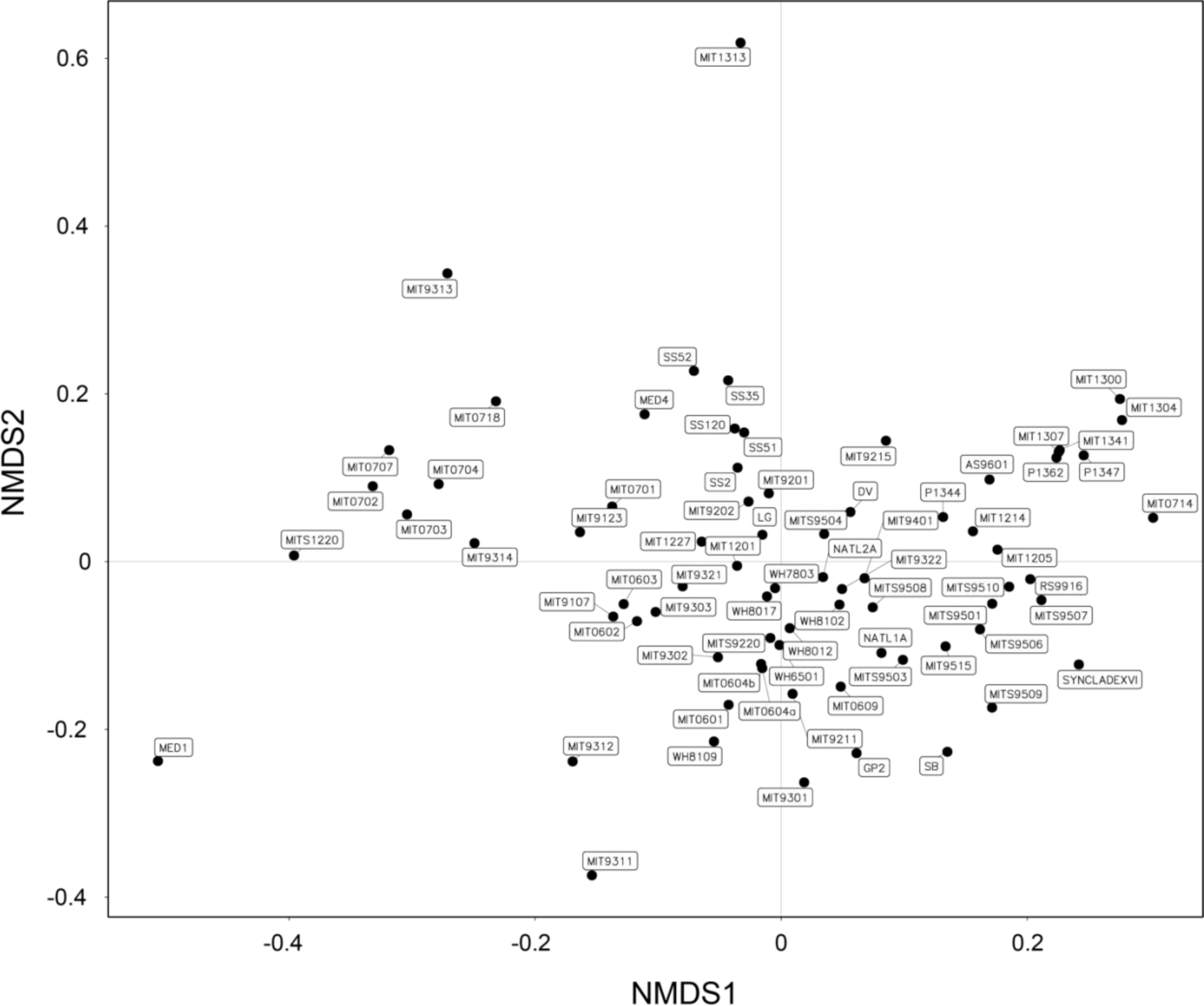
Mapping of cultures onto ordination plots labeled to indicate the position of each cyanobacterial host’s community at the ASV level in the ordination plots from Figure 4 and Figure S4. Non-metric multidimensional scaling (NMDS) of heterotroph communities using unweighted UniFrac as the distance metric. Each point in the NMDS plot represents a single heterotroph community associated with the given cyanobacterial host indicated in the name. The closeness of two points in NMDS space reflects the distance between communities, with communities having more similar phylogenetic structure (as measured by unweighted UniFrac) grouping more closely together.

**Figure S4.**
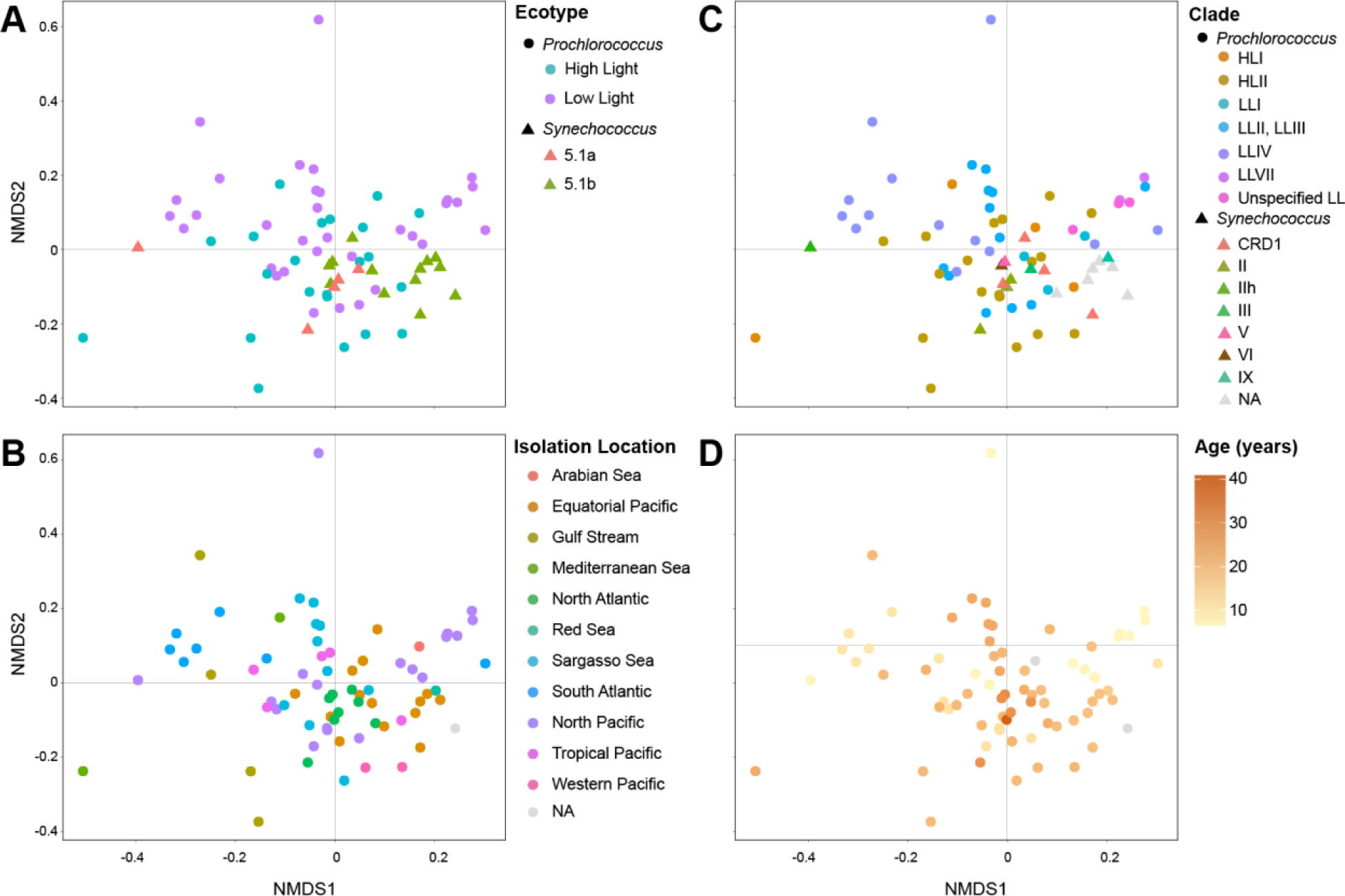
Ordination of heterotroph community composition at the ASV level overlaid with enrichment culture metadata. Non-metric multidimensional scaling (NMDS) of heterotroph communities using unweighted UniFrac as the distance metric overlaid with (A) cyanobacterial ecotype, (B) isolation location, (C) cyanobacterial clade, and (D) culture age. The closeness of two points in NMDS space reflects the distance between communities, with communities having more similar phylogenetic structure (as measured by unweighted UniFrac) grouping more closely together. Heterotroph communities for which metadata was not known are indicated as NAs. See also Figure 4, and Figures S3 and S7.

**Figure S5.**
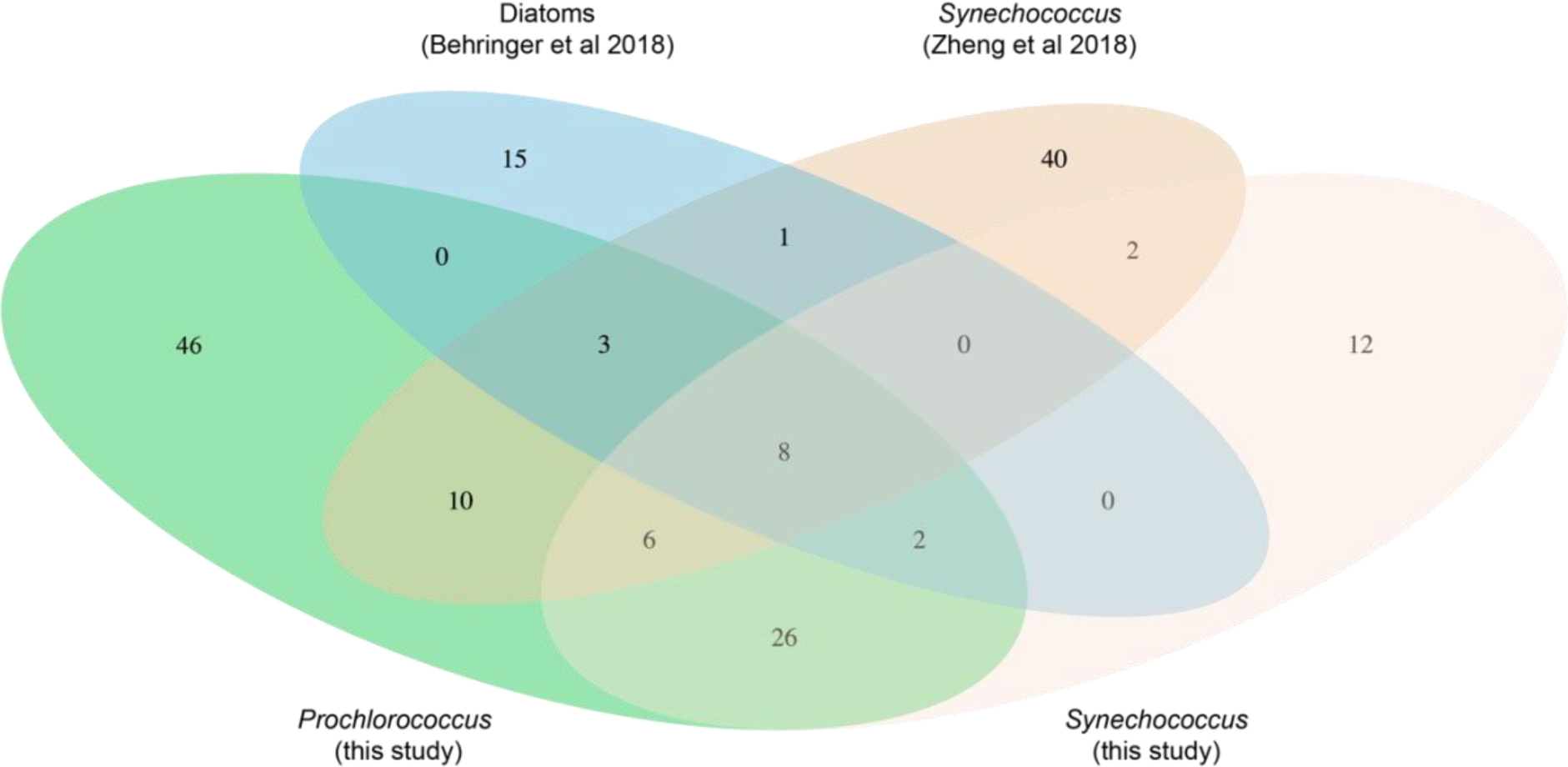
Venn diagram showing the number of OTUs shared in heterotroph communities from diatoms (Behringer et al., 2018) and *Synechococcus* (Zheng et al., 2018) compared to the *Prochlorococcus* and *Synechococcus* in this study. The area of the ellipses is not to scale with the number of OTUs.

**Figure S6:**
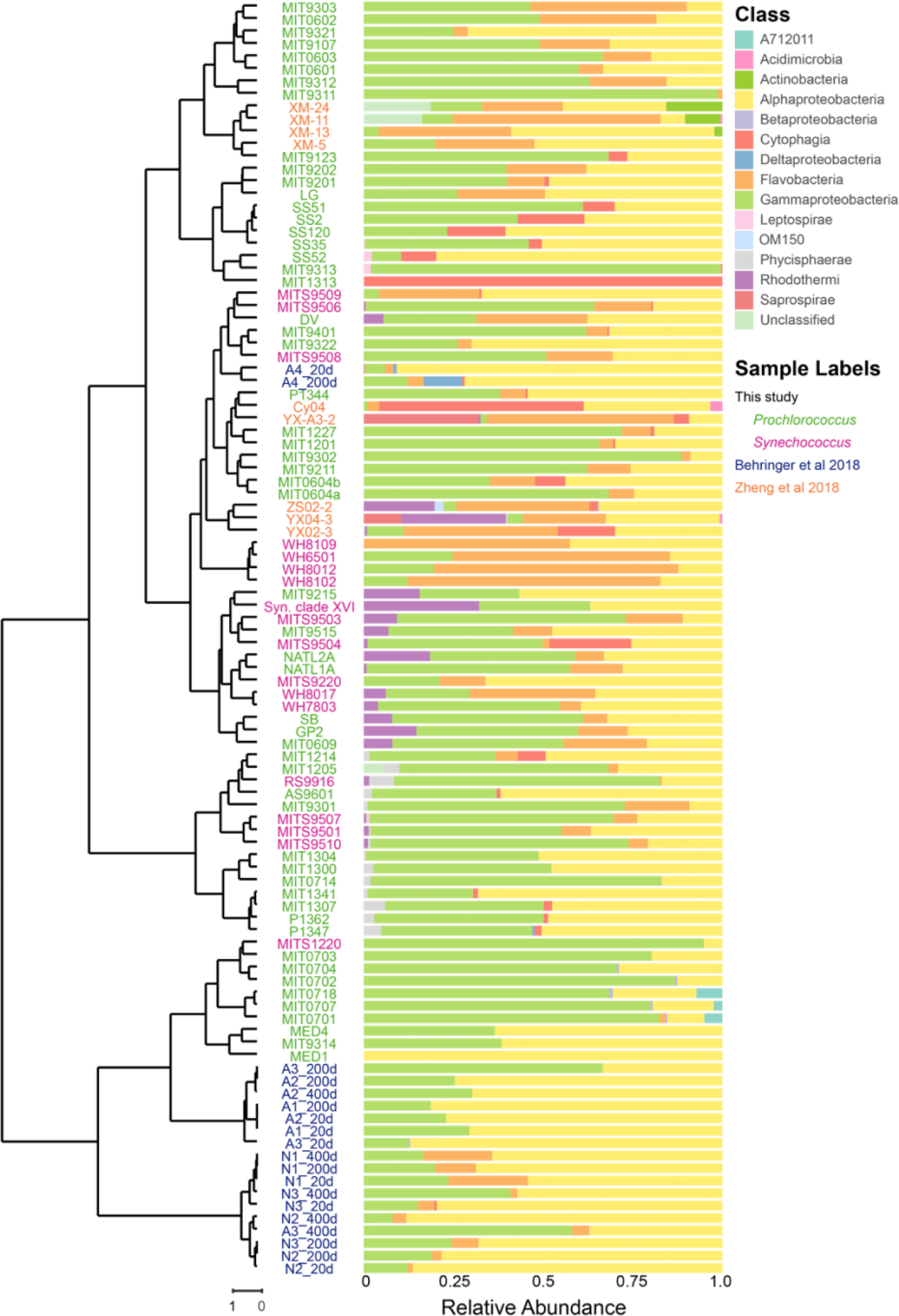
Heterotroph communities accompanying diatoms (dark blue) (Behringer et al., 2018) and *Synechococcus* (orange) in culture for less than a year (Zheng et al., 2018) compared to this study (green (*Prochlorococcus*) and magenta (*Synechococcus*)) with cultures over 5 years old. Bacterial classes are indicated by the colors in the legend, and relative abundance of each class in the heterotroph community of the corresponding culture is shown. Heterotroph communities in each cyanobacterial culture (vertical axis) are organized by the hierarchical clustering tree on the left using Ward’s method of hierarchical clustering with unweighted UniFrac as the distance metric on 97% OTUs. See also Figure 2 and Figure S2.

**Figure S7:**
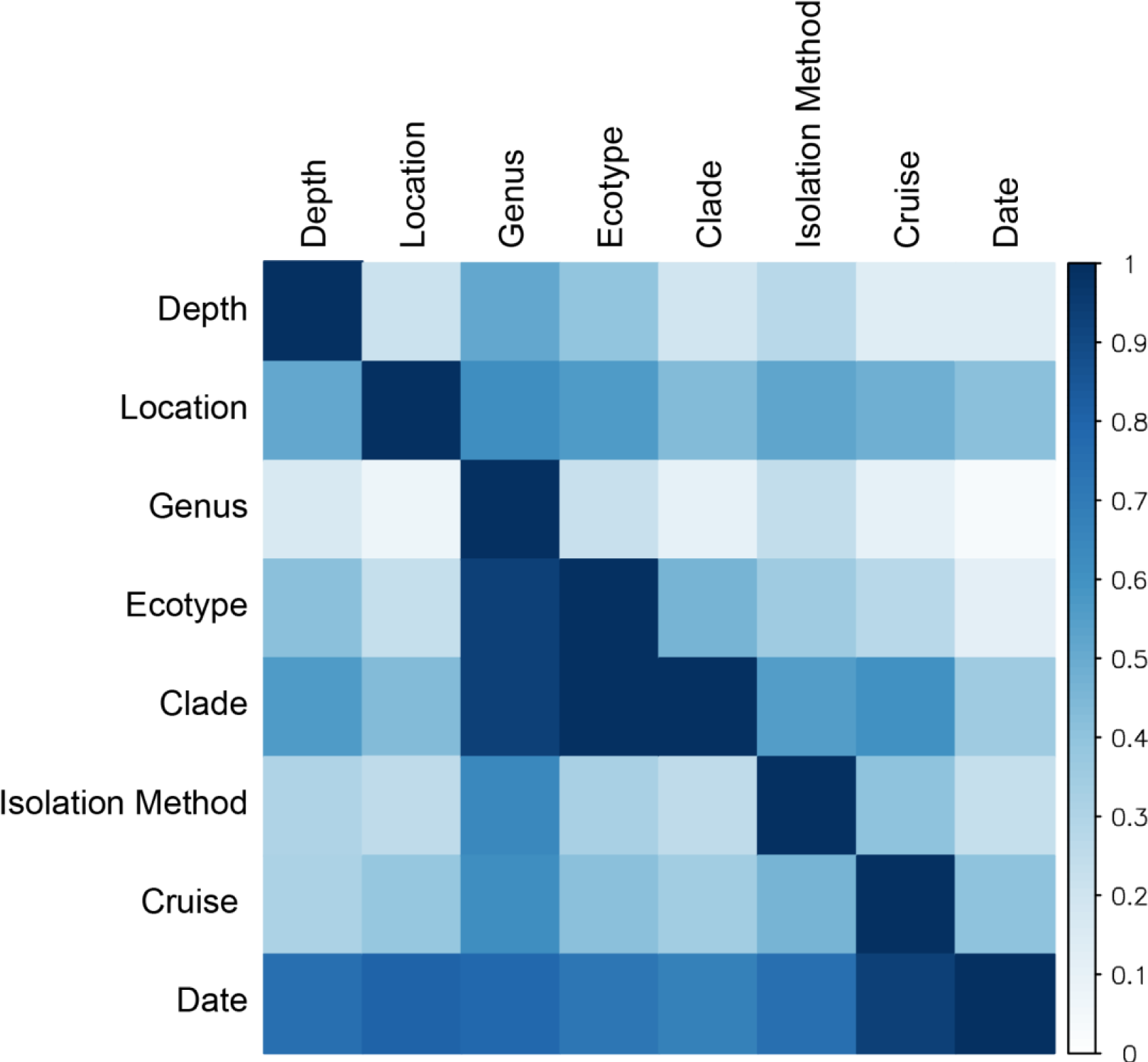
Associations between metadata pertaining to isolation or host phylogeny of enrichment cultures. The pairwise association between each variable indicated on the vertical and horizontal axes were calculated using an asymmetric measure of association, Goodman and Kruskal’s τ, ranging from 0 to 1. A τ of 1 indicates that there is a perfect correspondence between the levels in one variable and the record index. For instance, the clade has a τ of 1 with ecotype because clades are perfectly nested within ecotypes (that is, identification of the clade is sufficient to identify the ecotype). Because of the asymmetry in the metric, ecotype has a τ less than 1, but greater than 0 with the clade; ecotype only constrains identification of clades, but does not perfectly determine them. See also Figures 4, S3, and S4.

**Figure S8.**
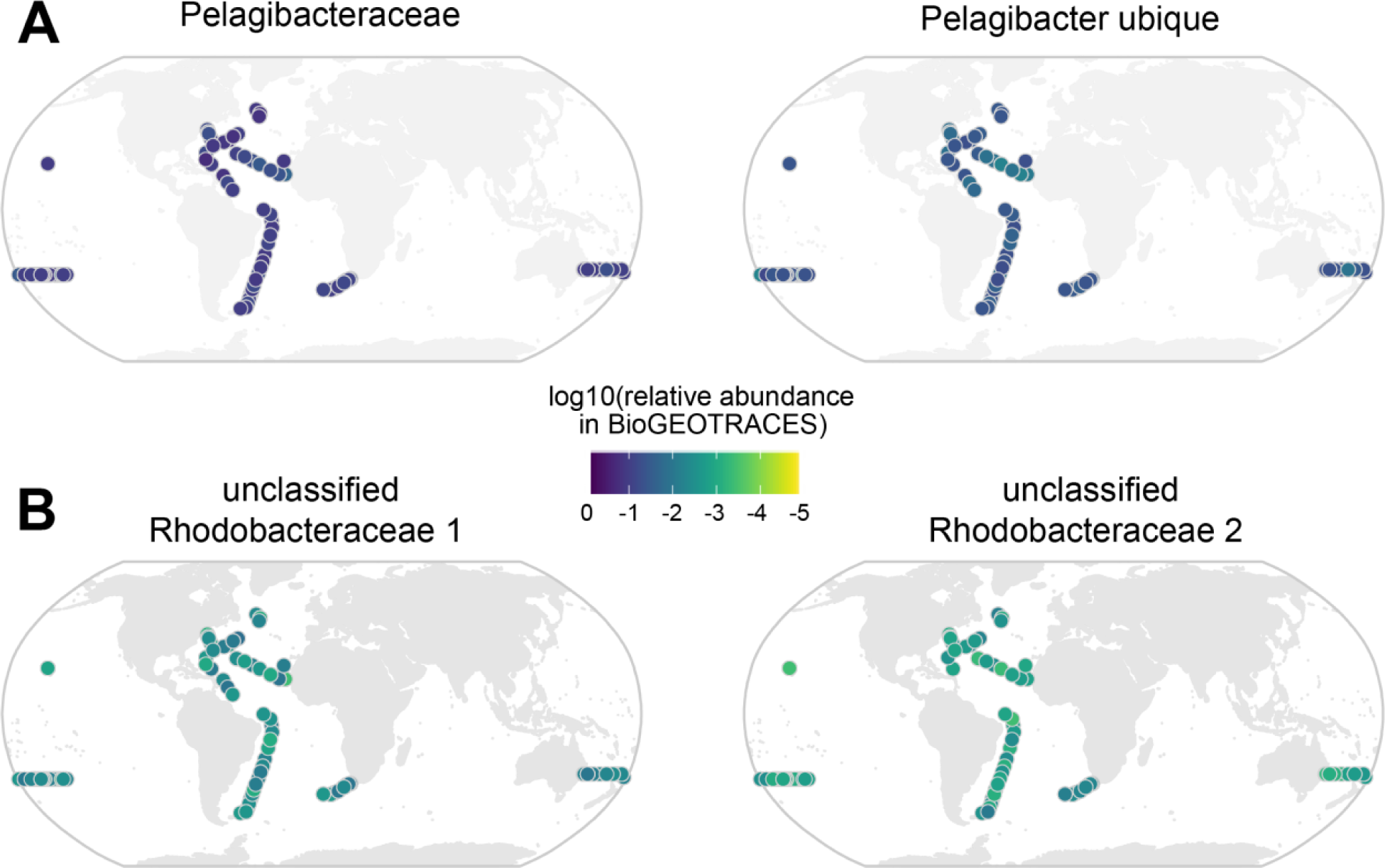
(A) Top two most abundant OTUs in bioGEOTRACES; these OTUs are not present in cyanobacterial cultures. (B) Top two most abundant OTUs (distinct unclassified OTUs in the family Rhodoobacteraceae) in bioGEOTRACES that are also present in the cyanobacterial cultures. Scale bar indicates the log10 relative abundance (number of reads normalized by number of non*-Prochlorococcus*, non-*Synechococcus* reads) at a given site. Related to Figure 5.

